# Probing the circuits of conscious perception with magnetophosphenes

**DOI:** 10.1101/449769

**Authors:** J. Modolo, M. Hassan, G. Ruffini, A. Legros

**Author notes:** Correspondance: Julien Modolo – - Laboratoire Traitement du, Signal et de l’Image (LTSI), Bâtiment 22 Campus de Beaulieu, 35042 Rennes Cedex, France. Co-first authors (equally contributed) (J. Modolo, M. Hassan).

## Abstract

**Background:** Conscious perception is thought to involve the large-scale, coordinated activation of distant brain regions, a process termed ignition in the Global Workspace Theory and integration in Integrated Information Theory, which are two of the major theories of consciousness.

**Methods:** Here, we provide evidence for this process in humans by combining a magnetically-induced phosphene perception task with electroencephalography. Functional cortical networks were identified and characterized using graph theory to quantify the impact of conscious perception on local (segregation) and distant (integration) processing.

**Results:** Conscious phosphene perception activated frequency-specific networks, each associated with a specific spatial scale of information processing. Integration increased within an alpha-band functional network, while segregation occurred in the beta band.

**Conclusions:** These results bring novel evidence for the functional role of distinct brain oscillations and confirm the key role of integration processes for conscious perception in humans.

## Introduction

The characterization of the physiological mechanisms that enable conscious perception is the focus of intense, multi-disciplinary research efforts. As opposed to the question of the nature of consciousness itself (referred to as the “hard problem” of consciousness (1)), the identification and characterization of the physiological processes that enable the emergence and maintenance of consciousness (referred to as the “soft problem” of consciousness (1)) is more readily achievable. Two theories of consciousness are currently dominant in the field, namely the Global Workspace Theory (GWT (2–4)), and the Integrated Information Theory (IIT (5)). While both share some key common points, such as information processing being at the core of conscious processes and that information needs to be *integrated* at a large scale to be efficiently shared and processed, IIT also posits that conscious experience involves a sufficiently “complex” level of information processing. Another recent theory, the Kolmogorov theory (KT) of consciousness (6), also establishes links between the algorithmic complexity of information processed by cortical networks and conscious processes. The idea that information processing coordinated between distant brain regions is essential has received considerable experimental support, notably from combined transcranial magnetic stimulation-electroencephalography (TMS-EEG) studies in disorders of consciousness (DOC) patients and healthy controls: during unconsciousness (e.g., deep sleep, Propofol anesthesia or coma), TMS-evoked EEG responses fail to propagate in cortico-cortical networks and only induce local activation, while complex spatiotemporal responses are evoked at the brain-scale during wakefulness (7).

One of the processes proposed to enable the large-scale cortical activation associated with conscious processes is termed *ignition* in GWT, and consists of a sudden and coordinated large-scale recruitment of distant brain regions that can make information available to the global workspace, enabling information processing (2). Ignition has been identified using functional MRI (fMRI) in a variety of tasks, as reviewed by Dehaene and Changeux (2). A compelling example is that, during subliminal perception of words, the activated brain network is considerably less spatially extended than during conscious perception (8). Using EEG and magnetoencephalography (MEG), it has also been shown that a large-scale activation in the beta band within the fronto-parieto-temporal network is involved in the report of target stimuli (9). Invasive intracranial recordings have also been performed in humans using deep electrodes during exposure to masked/unmasked words, which revealed an increase in gamma power in distant sites in association with a beta synchrony increase. In a recent experimental study, van Vugt et al. provided compelling evidence for the ignition process in monkeys, using intracranial electrodes during a phosphene perception task (10). Phosphenes are visual perceptions in the absence of light, and can triggered by electric stimulation (electro-phosphenes (11)) or magnetic stimulation (magneto-phosphenes (12)). In this study, van Vugt et al. performed intracranial stimulation of the visual cortex in monkeys at different current intensities (sub-threshold vs. supra-threshold) and linked visual saccades with phosphene perception. Consistent with predictions from GWT or IIT, large-scale brain activation was observed at the onset of phosphene perception.

In the present study, we aimed at inducing magnetophosphene percepts to characterize non-invasively the associated ignition process in humans. Our study involved twenty human subjects exposed to a 50 Hz magnetic field (MF) at 11 different flux density conditions given in a random order (5 mT steps ranging from 0 -sham-to 50 mT, each repeated 5 times for a total of 55 epochs per subject), while dense-EEG (64 channels) was recorded. Each epoch of exposure lasted 5 seconds, which was chosen to keep reasonable the duration of the experiment. MF exposure was delivered through a whole-brain exposure system designed and manufactured in-house (Fig. 1A), that exposed the whole brain and eyeballs to a homogeneous level of MF. Large-scale functional connectivity was estimated from dense-EEG measurements using source connectivity analysis (13), and graph theory metrics were used to quantify local and global/distant processing, enabling the identification of nodes in a cortical network. The study design is summarized in Fig. 1. Our hypothesis was that, during suprathreshold (conscious) magnetophosphene perception, a drastic increase in global/distant processing characteristic of ignition occurs, which should be detectable through an increase in network integration quantified by graph theory based metrics.

**Fig. 1.**
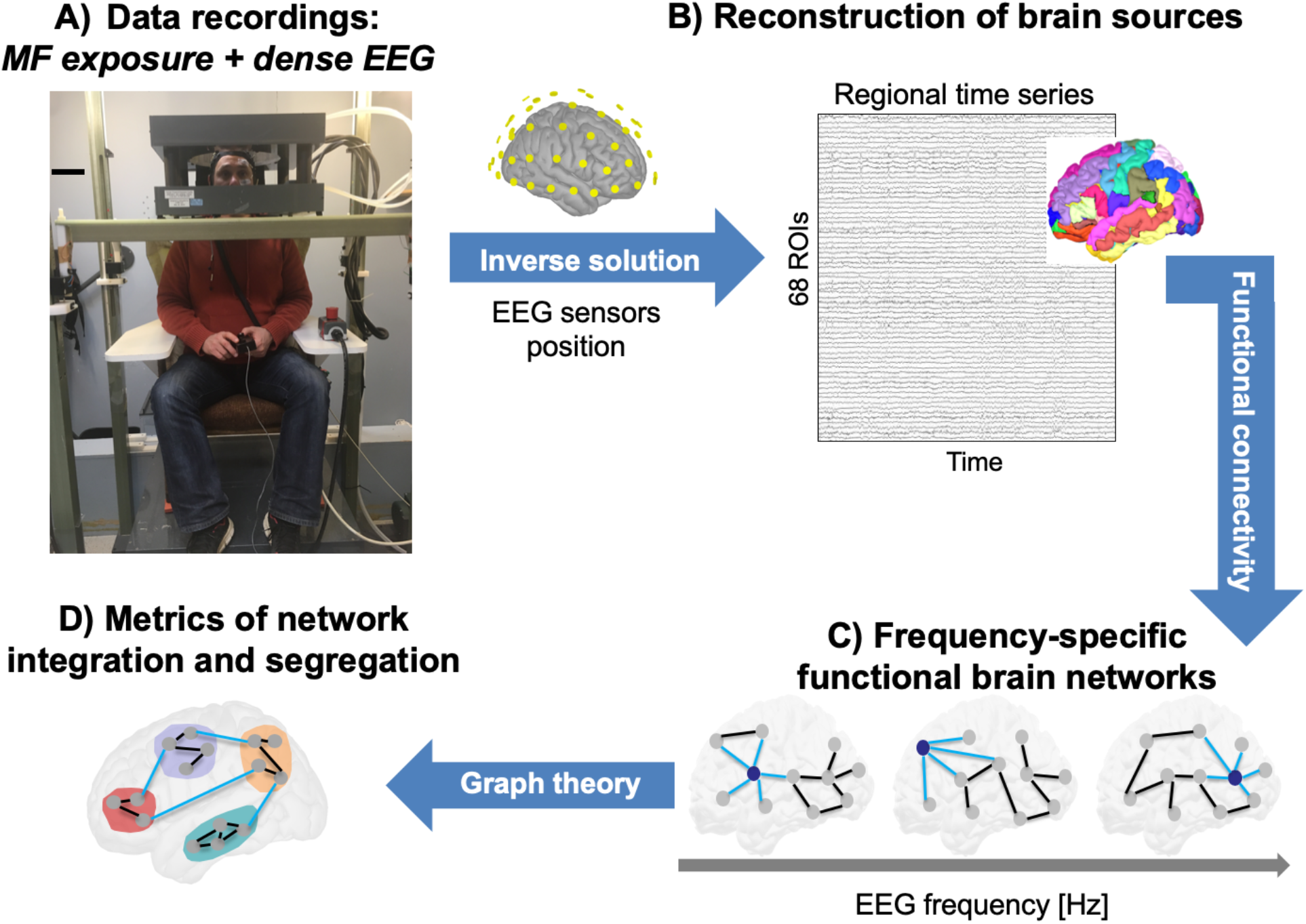
Study design and data analysis pipeline. **A)** MF exposure setup (water-cooled Helmholtz coils, in-house design and engineering, see Methods for details) used to induce magnetophosphene perception. Subjects could report magnetophosphene perception using a button press. Dense-EEG (64 channels, MRI compatible cap) was recorded throughout the entire experiment. **B)** Source activity was reconstructed for the 68 regions of the Desikan-Killiany atlas. **C)** Frequency-specific functional networks were computed for the theta, alpha and beta frequency bands. **D)** Metrics from graph theory were used to quantify segregation and integration (local and global processing, using clustering and participation coefficients, respectively).

## Materials and Methods

### Experimental protocol

This study was approved by the Health Sciences Research Ethics Board of Western University (London, ON, Canada) under the reference HSREB#18882. A total of N=20 healthy volunteers were recruited (age: 23.4 +/− 1.4, 10 males and 10 females). Exclusion criteria ensure that study participants had no cardiovascular, neurological or visual system disorder. They were equipped with a 64 MRI-compatible EEG cap (Maglink, Compumedics-Neuroscan, US). Subjects were asked to seat in the chair placed below the MF exposure device and the coils where lowered so subjects had their head centered in the device for the duration of the experiment. EEG was continuously recorded, and subjects were asked to remain eyes closed for the duration of the experiment. An adaptation time to the darkness of the room of 5 minutes was present at the beginning of the exposure sequence. The 55 MF exposure conditions were then randomly delivered through a custom-made LabView (National Instruments, USA) program, had a duration of 5 seconds each, and were separated by 5 seconds. Each time that the participant began to perceive magnetophosphenes, he/she had to press as fast as possible on a button-press directly synchronized with the EEG acquisition system.

### MF exposure device and physical principles

The MF exposure device was used to deliver whole-brain exposure and consisted of “Helmholtz-like” coils designed and manufactured in-house (for full details, please refer to (14)). Let us mention that Helmholtz coils are a standard setup in physics dating back to the end of the 19^th^ century. The mean diameter of each coils was 50 cm and their width 7.3 cm, and coils were separated by 20 cm. Water cooling (0.8 L/min) was used through the squared hollow wire in order to prevent coil heating. MRI gradient amplifiers (MTS corporation, USA) were used to power each coil (up to 200 A_rms_ capability). The experimental setup was Medical Grade approved by the Canadian Standard Association (CSA). The MRI gradient amplifiers driving a 50 Hz current through the coils, which per Maxwell-Ampere’s law generated a homogeneous 50 Hz MF at the level of the head and eyes, which resulted in an induced electric field at the same frequency in these exposed biological structures. Therefore, the MF exposure non-invasively generated induced fields and current in the brain and retinal tissues at 50 Hz. Furthermore, let us note that the homogeneity zone of the field included the entire brain (14).

### EEG acquisition and processing pipeline

64-channel EEG was acquired at a sampling frequency of 10 kHz and down-sampled to 1 kHz. EEG data was imported into Matlab, and band-pass filtered between (3-35 Hz) to remove artefacts present in the signal due to the MF exposure at 50 Hz. Importantly, the entire experiment was also performed using a phantom (watermelon) with the EEG cap that was setup in the exact same manner than with participants, to validate the pre-processing step. Two electrodes were connected to a signal generator providing a signal within the alpha range (10 Hz) and resulting in a signal recorded at the level of EEG electrodes in the same order of amplitude as in humans (approx. 50 µV). Using this ground truth for the recorded EEG signal, we verified that the reconstructed EEG signal after filtering of the 50 Hz exposure artifact was unaltered (not shown). For each EEG recording, the 55 EEG epochs (11 MF exposure conditions, each repeated 5 times) were extracted and imported in the Brainstorm Matlab package (15). The entire duration (5 seconds) of each epoch was extracted, and a window function was applied (Hamming window) to avoid edge effects.

Nasion-inion and preauricular anatomical measurements were made to locate each individual’s vertex site. Electrode impedances were kept below 10 kOhm. EEG signals are frequently contaminated by several artifacts sources, which were addressed using the same preprocessing steps as described in several previous studies dealing with EEG resting-state data (16, 17). Briefly, bad channels (signals that are either completely flat or contaminated by movement artifacts) were identified by visual inspection, complemented by the power spectral density. These bad channels were then recuperated using an interpolation procedure implemented in Brainstorm (15), using neighboring electrodes within a 5 cm radius. Epochs with voltage fluctuations > +80 µV and < −80 µV were removed.

### Magnetophosphene reports

Subjects were instructed to report magnetophosphene perception by using a button press as soon as they could when the perception began. Subjects were instructed to press only once, however a custom Matlab code was developed to detect and remove possible “double presses” which could accidentally occur.

### Brain networks construction

Functional brain networks were reconstructed using the EEG source-space connectivity method (13, 18), which includes two main steps: 1) reconstruct dynamics of the cortical sources by solving the inverse problem, and 2) measure the statistical couplings (functional connectivity) between the regional time series. The EEG source connectivity method links the recorded scalp EEG signals with the functional relationships between anatomical brain regions (e.g., networks), through the EEG inverse problem that provides the localization of the cortical sources originating these EEG signals. Several methods exist for both of these two steps (EEG inverse problem and functional connectivity measures). Here, we used the weighted minimum norm estimate (wMNE) algorithm as an inverse solution. The reconstructed regional time series were filtered in different frequency bands (theta, 4-7 Hz; alpha, 8-12 Hz; beta; 13-30 Hz). For each frequency band, functional connectivity was computed between the regional time series using the phase locking value (PLV) measure (19). The wMNE/PLV combination was chosen according to recent model-based and data-driven comparative studies of different inverse/connectivity combinations (20).

EEGs and structural MRI template (ICBM152) were co-registered through the identification of anatomical markers using Brainstorm (15). A Desikan-Killiany atlas-based segmentation approach was used, consisting in 68 cortical regions (21). The OpenMEEG (22) software was used to compute the three-layer (scalp, skull and brain) model. The functional connectivity between the 68 regional time series was computed using the PLV, using EEGNET (15, 23), for each condition, in the different frequency bands. This resulted for each subject, in each of the three frequency bands, in 55 68×68 connectivity matrices (one per MF condition). Finally, we only kept the strongest 10% of connections (proportional threshold).

### Characterization of functional networks

Our main objective was to explore two important properties related to information processing in the human brain network:

**- <u>Network segregation</u>**, which reflects local information processing. For this reason, the clustering coefficient ‘Cc’ was computed, considered as a direct measure of network segregation (24). In brief, Cc represents how close a node’s neighbors tend to cluster together (25). This coefficient is the proportion of connections among a node’s neighbors, divided by the number of connections that could possibly exist between them, which is 0 if no connections exist and 1 if all neighbors are connected.

**- <u>Network integration</u>**, which reflects global information processing. The participation coefficient was computed to measure the diversity of node inter-modular connections (26). The participation coefficient (Pc) of a node is close to 1 if its links are uniformly distributed among all the modules and 0 if all of its links are within its own module. Nodes with high participation coefficients interconnect multiple modules together, and hence can be seen as connectivity hubs. At the group level, this resulted in a Cc and Pc matrix for each of the 68 ROIs in each of the 55 MF conditions, for each frequency band.

### Statistical analysis

First, the Pearson correlation between the Cc/Pc matrix and the reported perception curve (probability of perception between 0 and 1, computed for each condition as the number of button presses divided by the number of repetitions, i.e. 5) was computed using Matlab and the False-Detection Rate (FDR) method was used to account for multiple comparisons. The graph theoretical metrics were computed using the Brain Connectivity Toolbox (27). Second, the linear trend associated with increasing MF flux density was assessed using the Jonckheere-Terpstra test (implemented in Matlab, The Mathworks, USA) with correction of the significance threshold using the FDR. The correlation with the (non-linear) perception curve was assessed using a linear correlation (Matlab, The Mathworks, USA) corrected using FDR.

### Quantification of the ignition process

To quantify the ignition process, the functional connectivity matrices were averaged over subjects at each flux density value (averaged also over five trials). The degree of each brain region, i.e. its number of functional connections, was then computed (after keeping the highest 50% of connections). The ignition process is therefore considered as the increased number of functional long-range connections when the probability of perception increases.

### Software

The functional connectivity, network measures and network visualization were performed using BCT (27), EEGNET (23) and BrainNet viewer (28), respectively.

The study design is summarized in Figure 1 below.

## Results

As expected, participants did not perceive phosphenes in the sham condition (0 mT), as seen in Fig. 2, and the perception of magnetophosphenes, as assessed by perception button-press self-reports, began with a 0.28 probability of perception at a threshold value of 20 mT when the entire head was stimulated by the homogeneous 50 Hz sinusoidal signal. It is worth noting that, at the maximal MF flux density value used (50 mT), the probability of magnetophosphene perception was close to 1 (0.96). Phosphene perception was reported by all subjects as colorless, stroboscopic dots/lines. The resulting magnetophosphene perception curve as a function of the MF flux density results in a sigmoid-like function, as can be seen in Fig. 2 (lower panel), similarly to the curve obtained in monkeys with increasing intracranial stimulation current values (10). The comparison between these two curves is direct: intracranial current injection results in an electric field that depolarizes neighboring neuronal elements, while MF exposure also induces an electric field in brain tissue per Maxwell-Ampere’s law of magnetic induction. One difference is that, in the present study, retinal cells instead of visual cortex cells are modulated by the induced electric field. As detailed in the Methods section, each functional network reconstructed in each of the 55 conditions featured 68 ROIs, and the participation and clustering coefficients quantifying *integration* (global) and *segregation* (local) processing respectively, were computed for each frequency band (theta, 4-7 Hz; alpha, 8-12 Hz; beta, 13-30 Hz). Gamma-band networks were not estimated due to the need of filtering the EEG from 50 Hz stimulation contamination, and to avoid possible biases. In the following, all *p*-values were corrected for multiple comparisons using the False Detection Rate (FDR) method.

**Fig. 2.**
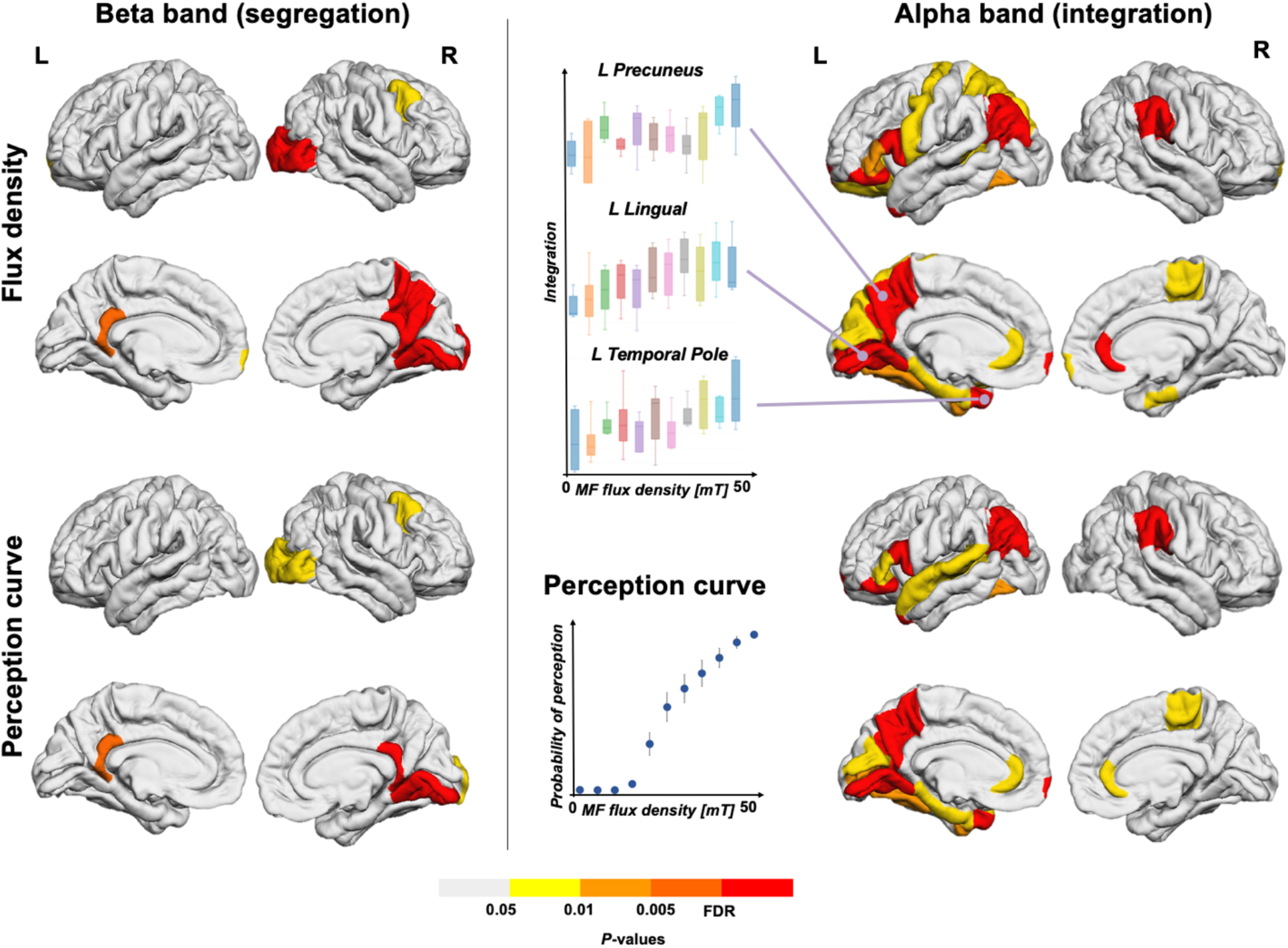
**Left panel.** Segregation results in the beta-band functional network (linear trend, flux density sub-panel; correlated with perception reports, perception curve sub-panel). Results are presented for each hemisphere. **Right panel.** Integration results in the alpha-band functional network (linear trend, flux density sub-panel; correlated with perception reports, perception curve sub-panel). Results are presented for each hemisphere. The increase in integration with flux density is shown for three regions of interest (left precuneus, left lingual and left temporal pole), along with the perception probability curve for magnetophosphene from subjective reports (N=20 subjects).

First, brain structures for which integration (quantified by the participation coefficient (29)) was correlated with the MF flux density were almost exclusively within the alpha band (see Fig. 2, right panel). More specifically, there were structures involved both in “subliminal” (i.e., 50 Hz MF exposure below the perception threshold) and suprathreshold magnetophosphene perception. These regions involved DMN (Default Mode Network) structures such as the right rostral anterior cingulate (*p*=0.0012) and left parahippocampal gyrus (*p*=0.0116); a structure from the VIS (visual) network (left fusiform, *p*=0.0089); and a DAN (Default Attention Network) structure (left pars triangularis, *p*=0.007). The left pericalcarine cortex (*p*=0.0052, visual cortex area), right enthorinal (*p*=0.0085), right frontal pole (0.0085), and right paracentral (*p*=0.0092) were also involved.

Regions correlating specifically with the perception curve involved a DMN region (left precuneus, *p*=0.001), two regions from the DAN (left pars orbitalis and left pars opercularis, *p*<0.001), and a region from the SAN (Salience Attention Network), namely the right supramarginal gyrus (*p*<0.001). Interestingly, the right supramarginal gyrus has been previously identified as involved in phosphene perception in humans using intracranial stimulation (30). The left temporal pole, *p* < 0.001, the left frontal pole, *p*=0.0026 and left inferior parietal, *p*<0.001 were also involved.

Second, magnetophosphene perception was associated with an increase in segregation (local processing) almost exclusively in a beta functional network, as illustrated in Fig. 2 (left panel). More specifically, the network segregation increased (both for “subliminal” and conscious perception) in the right lingual cortex (*p*<0.001, VIS), within DMN regions (right precuneus, *p*=0.0018; and left/right isthmus cingulate, *p*=0.0019 and *p*<0.001 respectively) and a DAN region (right caudal middle frontal gyrus, *p*=0.0031). In the case of conscious (suprathreshold) perception, only the right lingual cortex and right isthmus cingulate (*p* < 0.001) specifically showed significant correlation with the perception curve in terms of beta network segregation.

Notably, while magnetophosphene perception was mainly restricted to effects within alpha- and beta-band functional networks, a limited number of brain structures displayed increased integration within the theta (left pars triangularis, *p*<0.001; right middle temporal, *p*=0.001) and beta (left precuneus, *p*<0.001) bands. Furthermore, regarding network segregation, an effect was also identified in a single region (right superior frontal gyrus, *p*<0.001) of the alpha functional network. It should also be noted that there is an asymmetry between hemispheres in terms of regions involved. This is consistent, for example, with the known hemispheric specialization of occipito-temporal pathways depending on the spatial resolution of visual stimuli (31).

Finally, we deciphered the ignition process during the conscious perception of magnetophosphenes (Fig. 3) during increased MF flux density, corresponding to a higher level of retinal stimulation. In this example, the number of functional connections of the left precuneus (one of the regions achieving the highest significance regarding integration) to distant regions, including frontal regions, increases with the probability of perception.

**Fig. 3.**
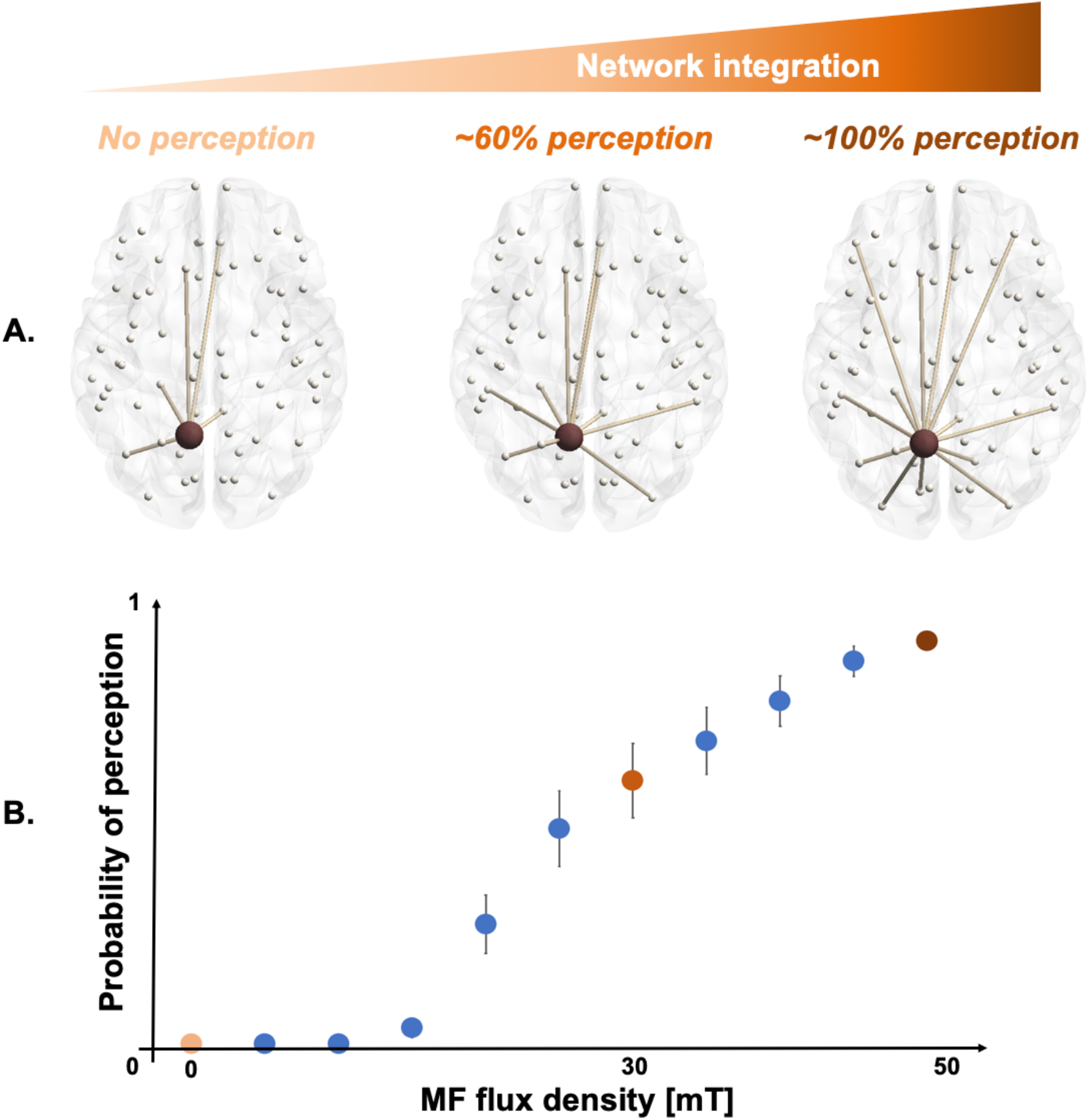
**A.** Illustration of the ignition process occurring with the increase of long-range connections (in this example, from the left precuneus) associated with the increase in probability of magnetophosphene perception at 0 mT (left), 30 mT (middle) and 50 mT (right). **B.** Probability of magnetophosphene perception curve from the N=20 subjects between 0 and 50 mT, with the 0 (light orange), 30 (orange) and 50 mT (burgundy) points highlighted.

This drastic increase in the functional connections between the left precuneus (a critical region from the DMN) and distant regions emphasize the central role of long-range connectivity for the emergence of conscious perception. Interestingly, long-range functional connections at baseline (without perception) remain present when the perception probability increases, while novel long-range connections gradually appear.

## Discussion

Magnetophosphene perception increases as a function of the increasing level of magnetic stimulation, which results from the induced electric fields modulating retinal neuron signaling (32). The increase in magnetophosphene perception probability induced in healthy human volunteers results in an increase of information integration at the brain-scale. Conscious perception was tracked using participant subjective reports and increased with gradually increasing stimulus strength. The increase of information integration was observed mostly in visual- and DMN-related regions, and also in a few frontal regions. We also identified an increase in the number of functional connections between one of the most critical nodes of the identified cortical network, namely the left precuneus, and distant regions, when the probability of perception increased. To our knowledge, this is the first use of EEG source-space networks to characterize the increase of integration information as a consequence of conscious visual perception. These results are in line with the recent study by van Vugt et al. (10) who identified the ignition process in monkeys, also using a phosphene perception paradigm (with stimulation delivered intracranially, as opposed to non-invasively in the present study). Taken together, these results provide further support for the ignition process that was conceptualized in GWT, and also for the integration concept at the core of IIT. Another originality of this study is that we characterized the functional networks involved in the biological effect occurring at the lowest known level of *in situ* electric field, which may provide further arguments for the use of magnetophosphene perception to define international safety electromagnetic exposure guidelines (33, 34).

One limitation in our study is that the whole brain was exposed to the delivered MF, which leaves open the possibility that some identified brain regions were not necessarily associated with phosphene perception, but rather with neuromodulation from the MF. Our whole head exposure protocol was chosen since it is the most effective to generate phosphenes perception non-invasively, which was the most important required feature at this initial stage of evaluating if brain networks would react to this perception. The MF exposure device was very different from a TMS (transcranial magnetic stimulation) device in the following respects: 1) the resulting MF flux density was much lower (0.05 T here versus > 1 T for TMS devices); 2) the level of dB/dt (proportional to the current induced in brain tissue) was much lower (approx. 15 T/s here versus > 10,000 T/s for TMS devices); 3) TMS targets a relatively narrow brain region typically at spiking suprathreshold levels, while the present exposure device provides whole-brain exposure at subthreshold levels (no neuronal firing outside of the retina can be directly induced by the MF used in this study even at the highest flux density −50 mT-, as detailed below); 4) TMS uses very short pulses (typically 100 µs) while the present MF exposure can deliver MF during minutes continuously. Next developments will use a more focal stimulus targeting the eyeball. That being said, one critical point is that the levels of MF flux density in this study are known to activate the retina only, since the perception of magnetophosphenes is the biological effect occurring at the lowest known MF flux density: it is precisely for this reason that the international guidelines protecting the general public and workers from the adverse effects from electromagnetic fields exposure are based, for extremely low-frequency fields (< 300 Hz), on the perception of magnetophosphenes (33, 34). Therefore, from the existing literature, direct acute effects of the MF delivered here on another system than the retina, is, according to the current existing literature, extremely unlikely. This supports further that the observed changes in functional connectivity are indeed associated with the perception of magnetophosphenes. Another limitation is that, since we used a 50 Hz stimulus, we were not able to investigate gamma-band functional networks. A 50 Hz frequency was selected because it is reported to be effective in triggering phosphenes perception (phosphene perception decreases as the stimulus frequency increases), although the related contamination of EEG signals involves filtering and thereby prevents investigation of gamma-related processes.

In addition to a contribution in terms of the fundamental mechanisms of consciousness, we also suggest that our experimental paradigm could form the basis of a novel protocol to probe conscious processes in humans. While a rather cumbersome experimental setup was used in this study, it would be possible to simplify it by using scalp (surface) electrodes placed on the temples and with an adapted level of delivered electrical current. Furthermore, this work identifies a key brain region in the conscious perception of a visual stimulus, namely the precuneus, a brain region that has been shown to be key for the maintenance of consciousness (35). Hence, our results bring support for a role of the precuneus not only in “basal” consciousness, but also in the perception of external stimuli.

Another more fundamental implication of these results is that the spatial extent of functional network dynamics and frequency processing is frequency dependent: during magnetophosphene perception, distant information processing increases mostly within the alpha band, while local information processing increases mostly within the beta band. This result is consistent with other studies pointing to the role of low-frequency oscillations for distant communication between brain regions, while higher frequency rhythms would rather contribute to stimulus encoding itself (36). Therefore, our results bring further support to the framework conceptualizing information flow and processing between brain structures as an interplay between slow and fast oscillations, each associated with a “range” for information processing and a specific functional role.

## Acknowledgments

### Funding

This study is funded by Electricité de France (EDF, France), Réseau de Transport d’Electricité (RTE, France), Hydro-Québec (QC, Canada), and in part by the Future Emerging Technologies Open Luminous project (H2020-FETOPEN-2014-2015-RIA under agreement No. 686764) as part of the European Union’s Horizon 2020 research and training program 2014–2018.

### Author contributions

Conceptualization: JM, AL. Data curation: JM, AL. Formal analysis: JM, MH, AL. Funding acquisition: AL. Investigation: JM, AL. Methodology: JM, MH, GR, AL. Project administration: AL. Resources: AL. Software: JM, MH. Supervision: JM, AL. Validation: AL. Visualization: JM, MH, GR, AL. Writing (original draft): JM. Writing (review and editing): MH, GR, AL.

### Data and materials availability

EEG data and analysis pipeline details are available upon reasonable request.

